# A simple solid media assay for detection of synergy between bacteriophages and antibiotics

**DOI:** 10.1101/2023.08.23.554535

**Authors:** Ethan Khong, Joseph Oh, Julian M. Jimenez, Roland Liu, Sage Dunham, Alisha Monsibais, Alison Rhoads, Pooja Ghatbale, Andrew Garcia, Ana Georgina Cobián Güemes, Alisha N. Blanc, Megan Chiu, Peiting Kuo, Marissa Proost, Ahnika Kline, Saima Aslam, Robert T. Schooley, Katrine Whiteson, Stephanie I. Fraley, David T. Pride

**Affiliations:** Department of Pathology, University of California San Diego, La Jolla, CA, USA; Department of Bioengineering, University of California San Diego, La Jolla, CA, USA; Department of Molecular Biology and Biochemistry, University of California Irvine, Irvine, CA, USA; Department of Medicine, University of California San Diego, La Jolla, CA, USA

**Author notes:** These authors contributed equally to this work (Ethan Khong, Joseph Oh).

**Keywords:** Synergy, Cooperativity, Antibiotics, Bacteriophages, Solid Media

## Abstract

The emergence of antibiotic resistant bacteria (ARB) has necessitated the development of alternative therapies to deal with this global threat. Bacteriophages (viruses that target bacteria) that kill ARB are one such alternative. While phages have been used clinically for decades with inconsistent results, a number of recent advances in phage selection, propagation and purification have enabled a reevaluation of their utility in contemporary clinical medicine. In most phage therapy cases, phages are administered in combination with antibiotics to ensure that patients receive the standard-of-care treatment. Some phages may work cooperatively with antibiotics to eradicate ARB, as often determined using non-standardized broth assays. We sought to develop a solid media-based assay to assess cooperativity between antibiotics and phages to offer a standardized platform for such testing. We modeled the interactions that occur between antibiotics and phages on solid medium to measure additive, antagonistic, and synergistic interactions. We then tested the method using different bacterial isolates, and identified a number of isolates where synergistic interactions were identified. These interactions were not dependent on the specific organism, phage family, or antibiotic used. *A priori* susceptibility to the antibiotic or the specific phage were not requirements to observe synergistic interactions. Our data also confirm the potential for the restoration of vancomycin to treat Vancomycin Resistant Enterococcus (VRE) when used in combination with phages. Solid media assays for the detection of cooperative interactions between antibiotics and phages can be an accessible technique adopted by clinical laboratories to evaluate antibiotic and phage choices in phage therapy.

## INTRODUCTION

The rise in antibiotic resistant bacteria (ARB) has become a global public health issue that threatens the lives of millions of people across the world every year (1). Among ARB, the ESKAPE pathogens (*Enterococcus faecium*, *Staphylococcus aureus*, *Klebsiella pneumoniae*, *Acinetobacter baumannii*, *Pseudomonas aeruginosa*, and *Enterobacter spp.*) are often multidrug resistant and are the leading cause of nosocomial infections. One potential solution to the growing threat of ARB is to use of bacteriophages (viruses that attack and kill bacteria) as alternative treatments to antibiotics. Phages thus far have largely been reserved for treatment of bacteria that are highly resistant to antibiotics (2) but could potentially have broader applications. There have been successful outcomes in a number of recent phage therapy cases (3).

Antibiotics are the current standard-of-care for the treatment of ARB infections. Since phages have not yet received regulatory approval, they are usually delivered in conjunction with antibiotics to ensure that the standard-of-care is met. When used in combination with antibiotics, it is difficult to determine the contributions of each to the eradication of the infection. In general, the field lacks randomized clinical trials to determine whether these combination therapies are effective (4, 5). One of the first steps towards determining whether these combination therapies can be effective is to investigate whether there are cooperative or even antagonistic interactions between bacteria and phage in *in vitro* systems. The lack of a standardized, accessible assay for determining cooperativity limits the field significantly.

The current methodology for determining whether there may be cooperative effects between antibiotics and phages is performed primarily in broth medium, where the target ARB is co-cultivated with antibiotic and phage. There are different methodologies to perform these broth assays (6-8), but no single procedure is universally accepted. Additionally, these assays are highly complex for clinical laboratory personnel, who need extensive training, and the assays require the acquisition of expensive equipment (9-11). While there have now been several studies to examine the cooperative phenomena between antibiotics and phages in broth (12), relatively little has been done to identify whether such relationships can be demonstrated on solid medium.

To address a growing need to understand the effects of the combination of antibiotics and bacteriophages against ARB, we sought to develop a cooperativity assay on solid medium. Such an assay can be performed without expensive equipment and has the potential to provide results that can be interpreted in a simplified fashion. Our goals were to: 1) develop an assay that can be easily performed in most clinical laboratories, 2) determine whether cooperative interactions between antibiotics and phages occur on solid medium for gram-positive and gram-negative bacteria, 3) decipher whether susceptibility to certain antibiotics and/or phages is necessary to demonstrate cooperativity, and 4) provide a template for straightforward interpretation of results without the need for complicated algorithms.

## METHODS

### Transient Diffusion in a Semi-Infinite Medium Approximation

A custom MATLAB (MathWorks, Inc) script was developed to model the diffusion of antimicrobial agents (antibiotic drug or bacteriophage) through an agarose medium. To model the perpendicular strips placed on an agarose plate, the concentration profiles of two agents diffusing perpendicular to each other were calculated. The semi-infinite approximation for diffusive mass transfer was used as previously described (13, 14) to predict the concentration of two agents: antibiotic (*α*) and bacteriophage (*β*), C_*α*_(x,t) and Cꞵ(y,t), as a function of distance and time (Eq. 1A, 1B) (**Tables S1 and S2**). The error function (Eq. 2A, 2B) and non-dimensionalized distance (Eq. 3A, 3B) were utilized to solve for the concentrations at each iterative distance and time interval. The following simplifying assumptions were made: 1-dimensional diffusion, dilute solution upon contact with agarose, transient diffusion. The concentration C_*α*_,source and C_*β*_,source *μ*g/mL were defined as an infinitely abundant sources C_*α*_(0,t) for x = 0 cm and C_*β*_(0,t) for y = 0 cm, respectively. The initial concentration C0 of all other points was defined as 0 *μ*g/mL for C_*α*_(x,0) and C_*β*_(y,0). By assuming that the agents diffuse a minute distance during the finite time of exposure relative to the size of the plate, we apply the semi-infinite medium approximation and set a boundary condition such that C_*α*_(∞,0) and C_*β*_(∞,0) = C_0_.

### Predicting drug interactions for equal concentration and equal diffusion coefficients

Using the semi-infinite medium approximation, contour plots of the concentration profile at different times were plotted on a 3 cm x 3 cm grid. Initially, agents *α* and *β* were modeled using equal source concentrations C_*α*,source_ = C_*β*,source_ = 1.0 *μ*g/mL and diffusion coefficient D_*α*_ = D_*β*_ = 1 x 10^-6^ cm^2^/s (15, 16) on the order of magnitude for an antibiotic drug diffusing through agarose. Different potential interactions between agents *α* and *β* were considered. No interaction between agents was modeled using the highest-single agent (HSA) model (17) (Eq. 4) (**Table S3**). This assumes that each agent acts independently, and the antibiotic effect of the combined agents is dictated by the higher concentration. Additive interactions were modeled following the assumption from the Loewe Additive Interaction model (17), i.e. concentrations of each individual agent can be added together as if they were the same agent (Eq. 5). Synergistic interactions are defined as interactions that result in a higher effect than an additive interaction (17). Synergistic interactions were modeled such that the effective concentration is the additive concentration plus the product of the concentrations, which is modulated by a coefficient k (Eq. 6). Antagonistic interactions are defined as interactions that result in a lower effect than the additive interaction (17). Antagonistic interactions were modeled using the assumption that each antibiotic agent is mutually antagonistic, with the overall effective concentration modulated by a coefficient q (Eq. 7). Minimum Bactericidal Concentration (MBC) curves were plotted over the concentration contours to visualize the resultant live bacterial lawn profiles.

### Predicting agent interactions for specific antibiotic drugs and bacteriophage combinations

Prediction for specific antibiotic (*α*) and bacteriophage (*β*) combinations were performed using the models above. However, parameters representative of the experimental conditions were used: C*α*,source = 1.5 *μ*g/mL and D_*α*_ = 1 x 10^-6^ cm^2^/s (15, 16) (for Vancomycin) and C_*β*,source_ = 1.2 x 10^-2^ *μ*g/mL and D_*β*_ = 5 x 10^-8^ cm^2^/s (18) (for phage Ben). Bacteriophage concentrations were converted from plaque forming units (PFU)/mL to *μ*g/mL by assuming that a PFU contains an average of 1 bacteriophage (19) and multiplying by the estimated molecular weight of an individual T4 bacteriophage, *e.g.* myovirus morphology, (20, 21) and converting to mass using Avogadro’s number (**Table S4**).

### Bacteria, phages, and culture conditions

Bacterial strains, including isolates of VRE, VSE, and STM were collected from the UCSD Center for Advanced Laboratory Medicine, under IRB#160524. All specimens collected were de-identified in such a manner that they could not be re-identified prior to their use in this study. Information, including antibiotic susceptibilities and speciation for each microbe was recorded (**Tables S5 and S6**). All isolates were identified to the species level using Maldi-Tof (Brucker, Billerica, MA), and antimicrobial susceptibilities using microbroth dilution on the BD Phoenix using panels PMIC-107 for gram positives and NMIC-307 for gram negatives. All the strains of bacteria and phages were cultivated in liquid Brain Heart Infusion (BHI) medium at 37°C with shaking at 250 rpm. BHI plates were made with an equal volume of 20 ml of BHI broth infused with 1.5% agar. All phages used in this study were isolated and purified from environmental sources using multiple enrichment protocols as previously described (22).

### Preparation of antibiotic and phage strips

Grade 1 Whatman filter paper (VWR, Visalia, CA; CAT no. 1001-150) was used to make a 5mm x 28mm paper strip cut out using Cricut Explore Air 2™. The strips were autoclaved, and then soaked in prepared antibiotic stock solutions matching the antibiotic concentrations of the standard antibiotic disks. For example, for vancomycin, the strips were soaked in 1.4 mg/µL concentration of a vancomycin stock solution. Standard antibiotic stock concentrations used in this study are listed (**Table S7**). Similarly, phage strips were prepared using high titer (10^8^ pfu/ml) phage stock. All the antibiotic and phage strips were soaked in their corresponding solutions for 12 hours at 4°C. Strips were dried in a biosafety cabinet for 1 hour without light exposure. Dried strips were used within 1 hour of drying or were stored at 4°C for up to 12 hours before use.

### Plating, stamping, and interpretation

For each isolate, an overnight culture was diluted to 0.2 OD_600_ and incubated for 15 min with shaking at 37°C. Then, 100uL of culture was combined with 3 mL of warmed 0.3% top BHI agar (BHI broth with 0.3% agar) and poured over 1.5 % BHI agar plate evenly to make a bacterial lawn. The plates with bacterial lawn were dried for two hours at room temperature before performing a stamping procedure with both phage and antibiotic strips aligned at a 90-degree angle on a predesigned L-shape stamp (**Figure S4**). Dried plates with the bacterial lawns were then inverted with the cover off and gently lowered on the L-shape stamps until top agar pressed minimally against the aligned strips. Then, the stamped plates facing upward were incubated at 37°C with no shaking for 18-20 hours. Control plates were prepared in a similar manner using the same stocks of antibiotic- or phage-impregnated strips. The control plates also contained blank autoclaved strips along with antibiotic disks (at the concentrations specified in **Table S7**), and a 4uL spot of liquid phage stock placed directly onto the agar plate. After 18-20 hours of incubation, plates were then imaged and analyzed.

## RESULTS

### Development of a solid media phage/antibiotic cooperativity assay

Our solid medium cooperativity assay design is based on the principle of impregnating separate filter paper strips with antibiotics and phages, placing the strips at a right angle on a lawn of bacteria, and then measuring growth inhibition along each strip (**Figure 1 and Figure S1**). If there is evidence of cooperativity between the phage and the antibiotic, a zone of cooperativity will form at the right angle created by the antibiotic- and phage-impregnated strips (**Figure S2**).

**Figure 1:**
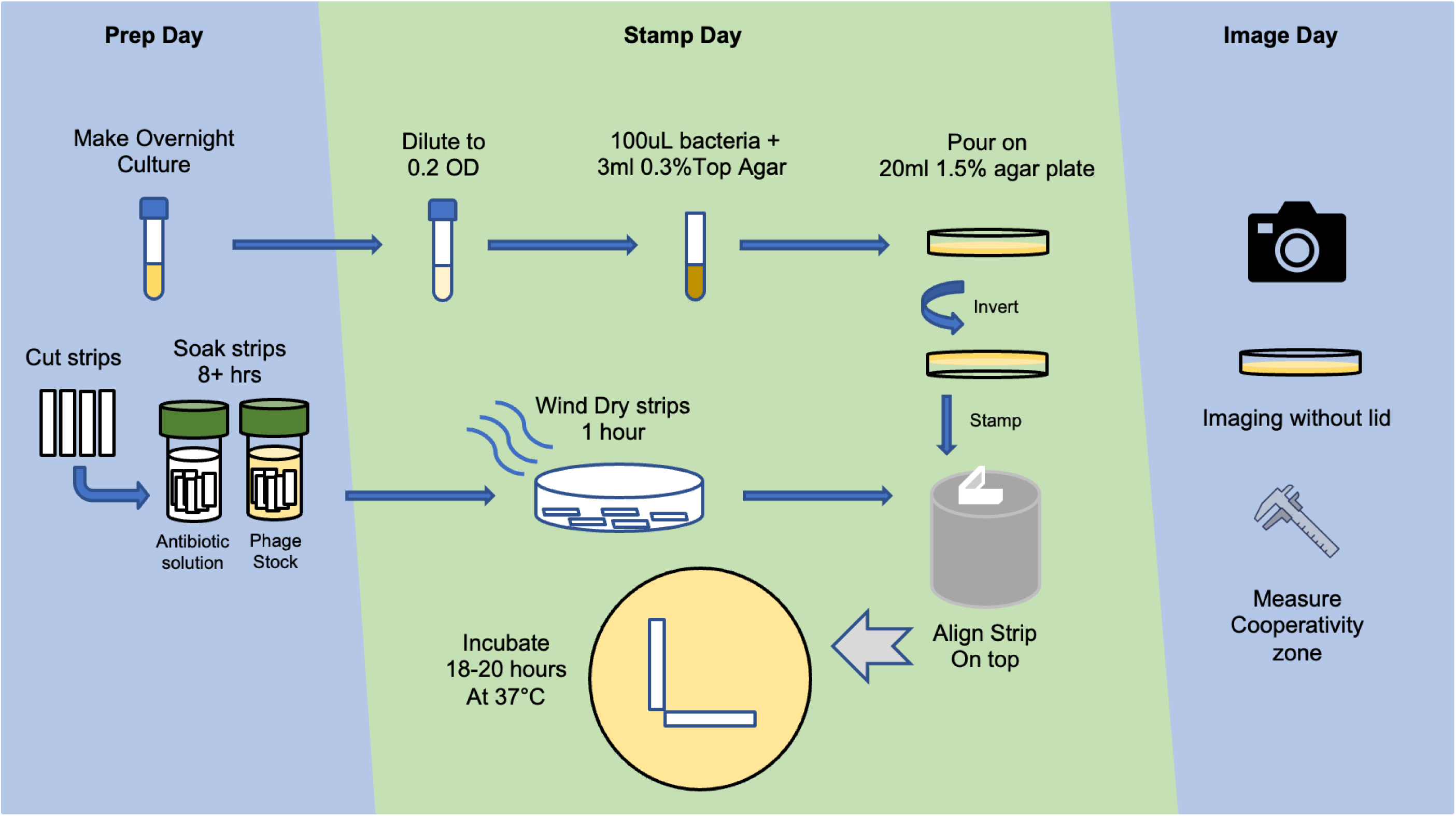
Workflow for phage-antibiotic cooperativity assays.

### Measuring additivity, cooperativity, and antagonism

We developed a custom model to predict the effective concentrations of antibiotics and phages as they diffuse away from their source strips through the agar medium and interact with the bacterial lawn. We did so using the semi-infinite medium approximation for unsteady-state mass transfer (13, 14), which predicts the concentration profiles of the antibiotic and phage as a function of distance from the strips and time (**Table S1**). We assumed that the depth of the medium was negligible compared to the width and only modeled diffusion in the top plane of view of the plate, setting boundary conditions and parameter values for the semi-infinite medium approximation based on a combination of measured, estimated, and literature values (**Table S2**).

We developed this model based upon the concept that we could observe killing of the bacterial lawn in areas that diffuse from the antibiotic or phage impregnated strips, which would reflect an effective minimum bactericidal concentration (MBC) (**Figure S3**). The interface between live and dead bacteria would create a profile that aligns with the MBC that is achieved by the combinatory antibiotic and phage effect. We then could develop a computational model to predict the concentrations of the antibiotic and phages as they diffuse across the agar using contour plots that represent different experimental results (**Figure 2**). Model parameters representative of the experimental conditions (**Table S2**) were used to predict the bacterial lawn profile under different antibiotic and phage interactions (**Table S3**) after 20 hours of incubation assuming an MBC of 0.1 *μ*g/mL. k and q are tunable variables that represent different extents of synergistic or antagonistic interactions (**Table S2**). We performed such simulations using a gram-positive model organism, *Enterococcus* spp. and several different Enterococcus phages (**Table S4**). Initial concentration and coefficients of diffusion representative of Vancomycin (C*α* =1.5 *μ*g/mL D*α* = 1 x 10-6 cm2/s) and Enterococcus phage Ben (C*β* =1.2 x 10-2 *μ*g/mL D*β* = 5 x 10-8 cm2/s) were used. We modeled no interaction (**Figure 2, Panel A**), and additive interactions between antibiotic and phage (**Panel B**). Our models displayed distinct convex curvatures that were indicative of strongly synergistic interactions (**Panels C and D**). For example, the model has different convex curvatures based on the extent of synergy displayed, with at least 1e6 greater killing (**Panel C**) or 1e12 greater killing (**Panel D**). We also could model antagonistic interactions between antibiotics and phage, which demonstrated concave curvatures (**Panel E**).

**Figure 2.**
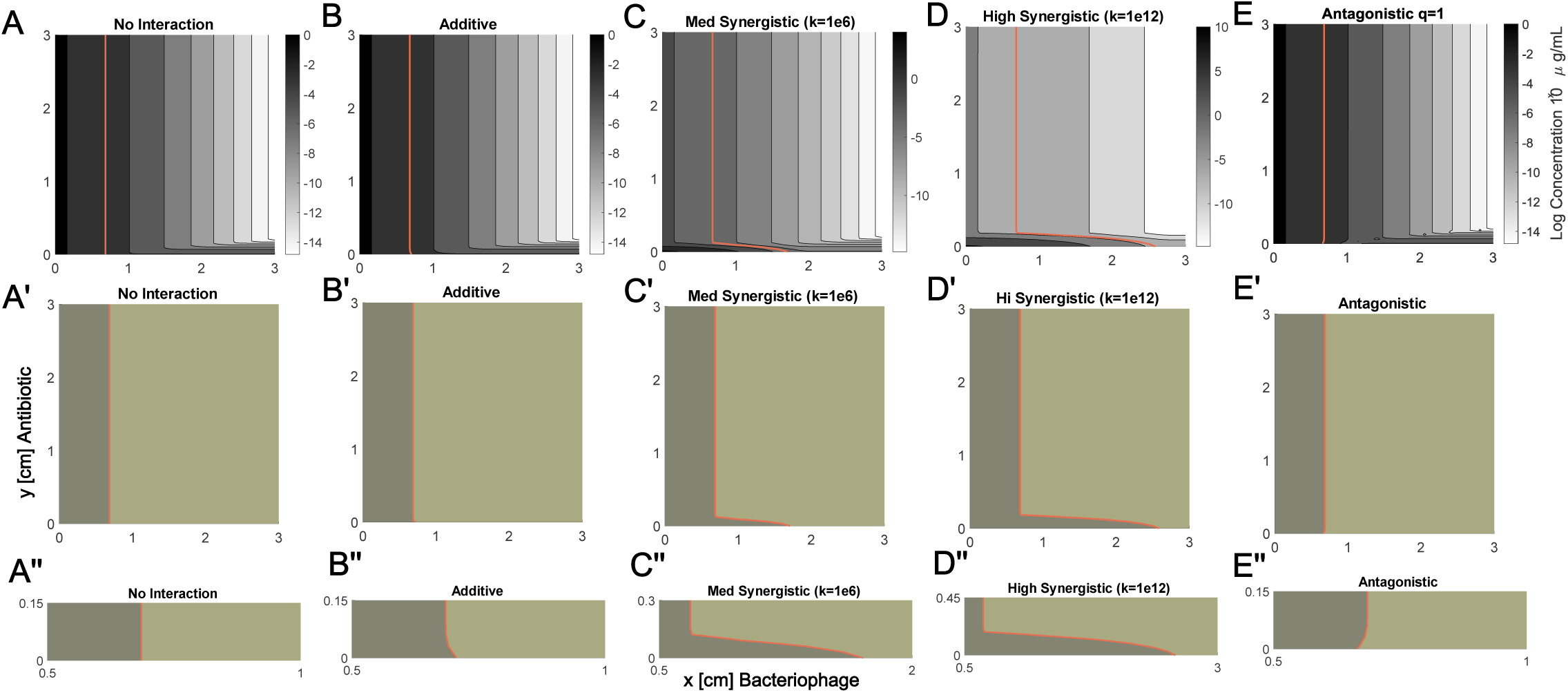
Model with parameters predictive of experimental results. Prediction of antibiotic (*e.g.* vancomycin) and bacteriophage (*e.g.* Ben) profiles based on different potential interactions. Concentration contour plots for representative antibiotic (C_*α*_ =1.5 *μ*g/mL D_*α*_ = 1 x 10^-6^ cm^2^/s) and bacteriophage (C_*β*_ =1.2 x 10^-2^ *μ*g/mL D_*β*_ = 5 x 10^-8^ cm^2^/s). (A) No interaction (B) Additive (C) Synergistic “medium” k = 1e6 (D) Synergistic “high” k = 1e12. (E) Antagonistic q = 1. Assuming MBC = 0.1 *μ*g/mL (red). Panels A’-E’ and A”-E” represent magnifications of portions of the panels shown in panels A-E, respectively.

### Evaluation of cooperativity in Vancomycin Resistant Enterococcus (VRE)

We next set up this solid media cooperativity assay to determine whether we could observe patterns similar to those predicted in the model (**Figure 2**). We expected to observe additional killing at the right angle where concentrations of the phage and antibiotic may be below the MBC of each individual phage or antibiotic, but together show cooperativity (**Figure S2**). A separate stamping device/procedure was developed to allow for the placement of the antibiotic and phage strips at perfect right angles on the medium (**Figure S4**). Each experiment was performed in triplicate to verify the accuracy and reproducibility of the results (**Figure 3**).

**Figure 3.**
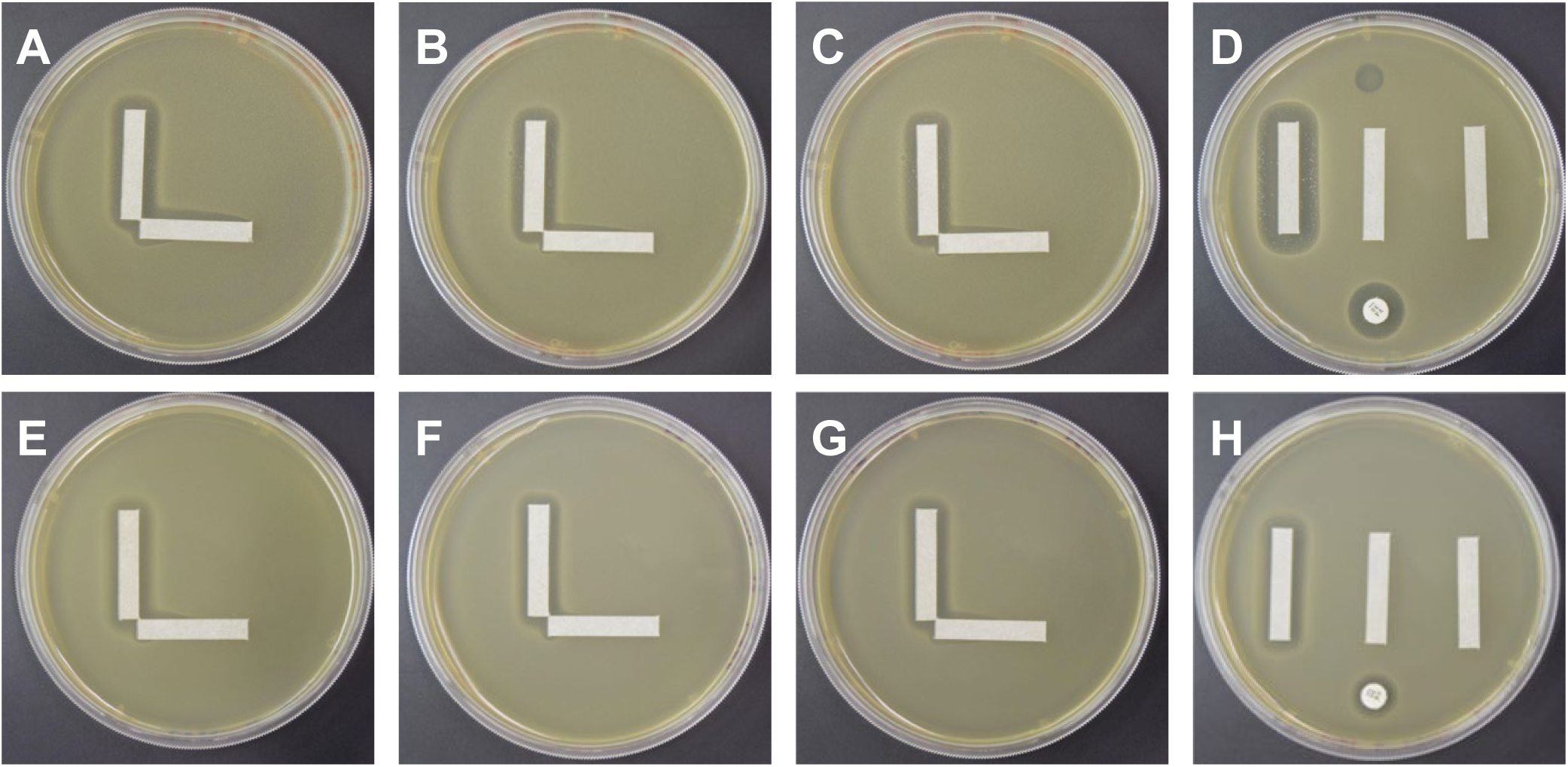
Solid media cooperativity assays for Vancomycin Resistant Enterococcus (VRE). Each specimen was tested with vancomycin (vertical strip) and a bacteriophage (horizontal strip). *E. faecium* EF98PII (VRE) with phage Bop is demonstrated in Panels A-D, where A-C represent 3 separate replicates of the cooperativity assay, and panel D represents the control plate with a vertical vancomycin strip (left), blank strip (middle), and phage strip (right), antibiotic disk (bottom), and phage spot (top). *E. faecalis* V587 (VRE) with phage Bop is demonstrated in Panels E-H, where Panels E-G represent separate replicates and Panel H represents the control plate.

We performed solid media cooperativity experiments for isolates of both *Enterococcus faecium* and *Enterococcus faecalis* (**Table S5**). We chose a set of phages that were active against the Enterococcus isolates (**Table S8;** genomes and further information about phage sources are available in (23). These isolates (for both species) may become resistant to vancomycin through expression of genes for enzymes that alter cell wall amino acid composition, often contained on a plasmid (24). We first used the cooperativity assay to examine a highly antibiotic resistant VRE isolate of *E. faecium* (EF98PII). We set the assay up with vancomycin as the antibiotic and Bop (myovirus) as the phage (**Figure 3**). While EF98PII is susceptible to Bop, it does not demonstrate complete lysis (**Panel D**). There is significant evidence in each of the replicates of a cooperativity zone between vancomycin and the phage (**Panels A-C**). We also identified similar interactions when *E. faecalis* was used rather than *E. faecium*, indicating that the cooperativity in VRE is not a species-specific phenomenon (**Figure 3, Panels E-H**).

We further examined the synergistic interactions observed for vancomycin and phage Bop for the *E. faecium* and *E. faecalis* VRE isolates (**Figure 3**). By measuring the extension of the zone of inhibition for *E. faecium* EF98PII, we were able to estimate the synergy coefficient (“k”) for vancomycin and phage Bop. Our results indicate that k = 1e6 (**Figure 4, Panel A**), which matched our model for medium level synergistic interactions between the phage and antibiotic. For *E. faecalis* V587, the coefficient was 1e16 for vancomycin and Bop (**Panel B**), indicating that high level synergy was observed. These data confirm that synergistic interactions occur between the antibiotic vancomycin and phage Bop for both *E. faecium* and *E. faecalis* isolates.

**Figure 4.**
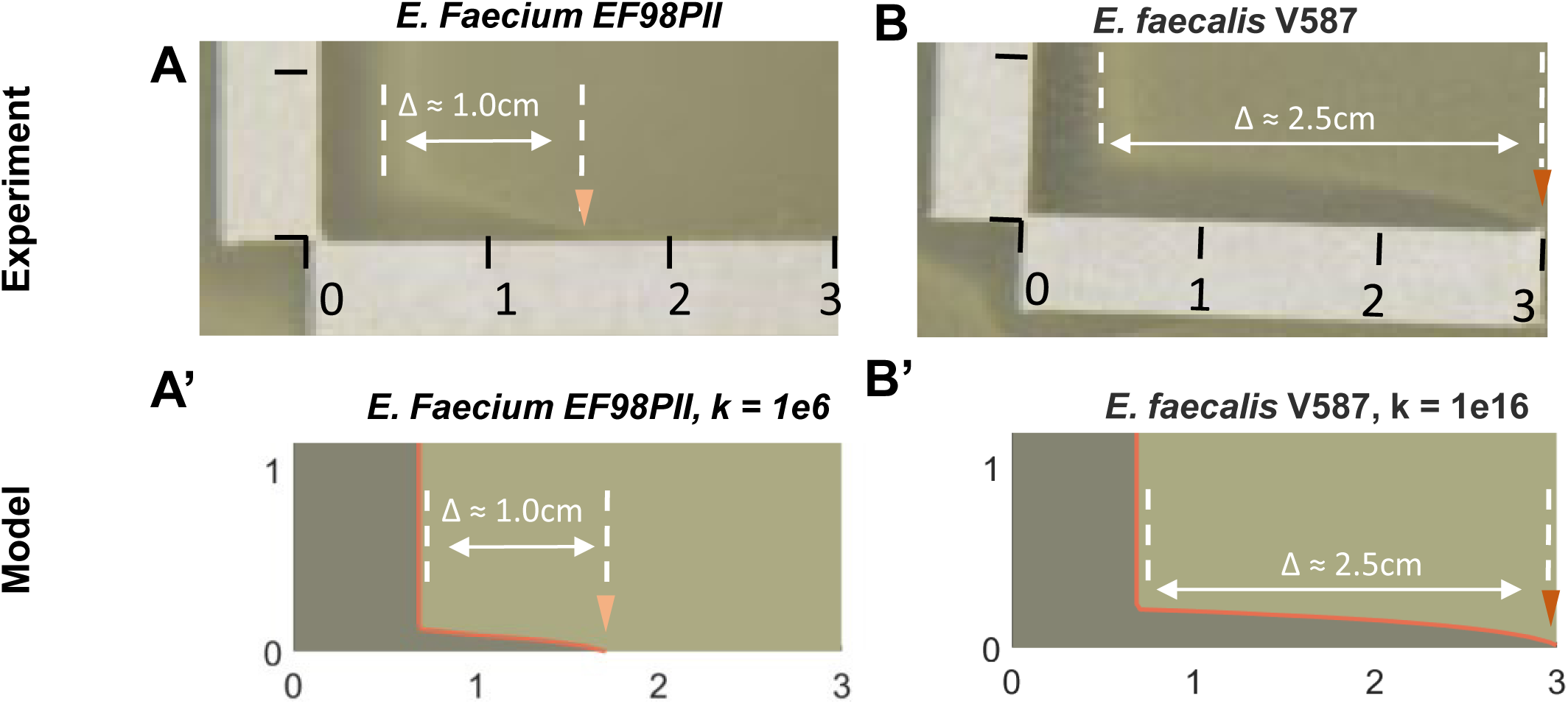
Comparison of experimental results and model predictions. (A) *E. faecium* EF98PII (VRE) treated with vancomycin (vertical strip) and bacteriophage Bop (horizontal strip). This resulted in a synergistic profile that extended 1.0 cm from the leading edge of the vertical zone of inhibition. (B) *E. faecalis* V587 (VRE) treated with vancomycin (vertical strip) and bacteriophage Bop (horizontal strip). This resulted in a synergistic profile that extended 2.5 cm from the leading edge of the vertical zone of inhibition. Model predictions for *E. faecium* EF98PII (A’) and *E. faecalis* V587 (B’) showed similar synergistic profile extensions and dimensions when the synergy coefficient was adjusted from medium synergy (k=1e6) to high synergy (k=1e16).

We also evaluated whether a 2nd class of antibiotics against VRE isolates demonstrated cooperativity with phages. We used *E. faecium* NYU in combination with linezolid and phage Bob (myovirus). In each of the replicates, we identified interactions that matched the synergy model (**Figure 5, Panels A-D**). We identified similar results for *E. faecalis* B3286 with phage PL (siphovirus), indicating that multiple different Enterococcus species can demonstrate similar results even with different phages.

**Figure 5.**
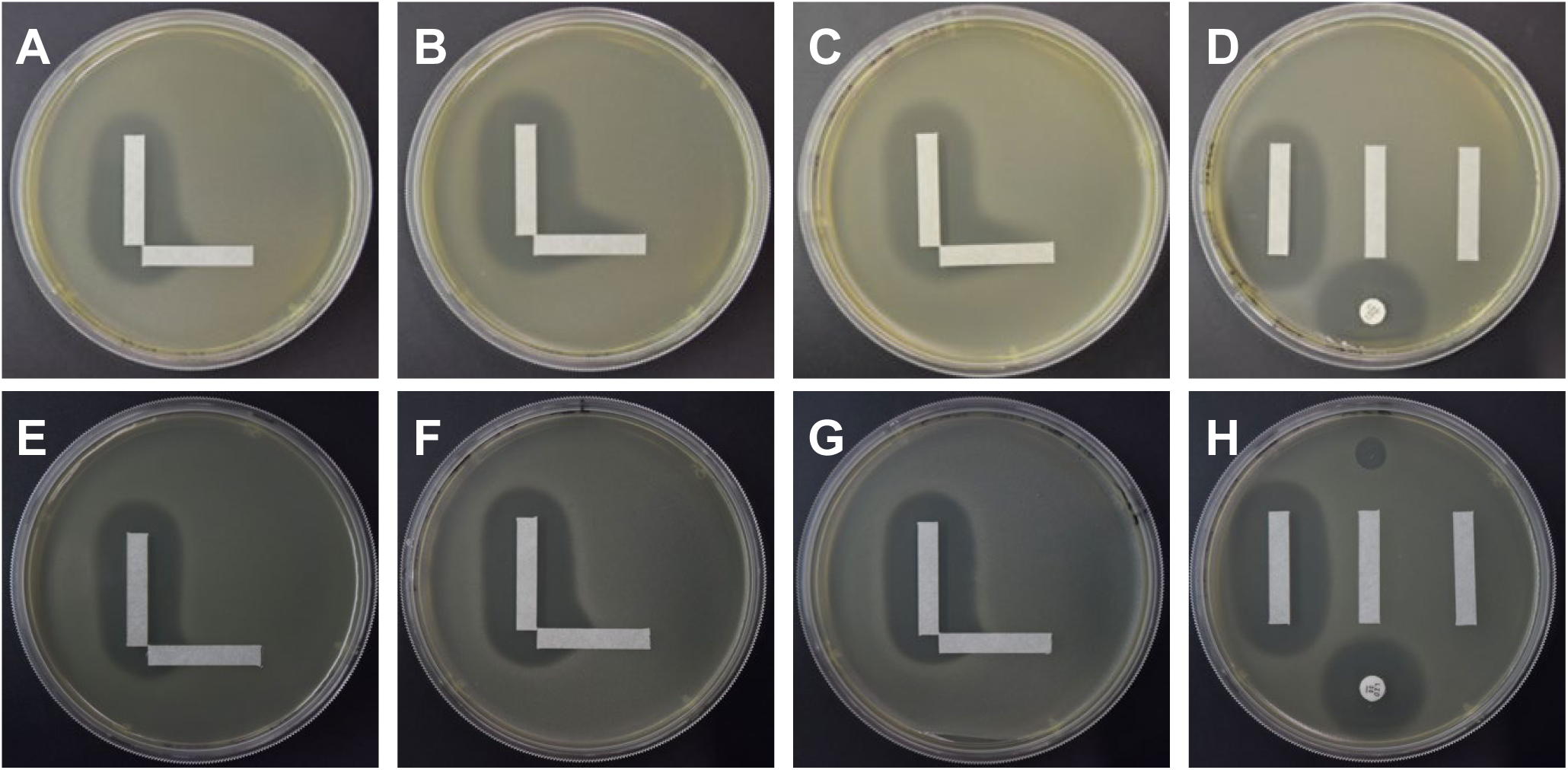
Solid media cooperativity assays for Vancomycin Resistant Enterococcus (VRE). Each specimen was tested with linezolid (vertical strip) and a bacteriophage (horizontal strip). *E. faecium* NYU with phage Bob is demonstrated in Panels A-D, where A-C represent 3 separate replicates of the cooperativity assay, and panel D represents the control plate with a vertical linezolid strip (left), blank strip (middle), and phage strip (right), antibiotic disk (bottom), and phage spot (top). *E. faecalis* B3286 with phage PL is demonstrated in Panels E-H, where Panels E-G represent separate replicates and Panel H represents the control plate.

We also performed the same cooperativity assay with a beta lactam antibiotic. Because *E. faecium* is intrinsically resistant to most beta-lactam antibiotics, we performed this assay using ampicillin along with phage Bob (myovirus). We also observed a significant interaction at the intersection of the antibiotic and phage indicating the presence of synergy (**Figure 6, Panels A-D**). These data suggest that while *E. faecium* isolates are resistant to certain antibiotics, the combination of these antibiotics with phages can lead to much greater killing. *E. faecalis* often is not resistant to beta lactam antibiotics such as ampicillin. We also noted significant synergistic interactions with phage Bop (myovirus) was used in combination with ampicillin (**Panels E-H**). These data suggest that there may be common mechanisms that lead to antibiotic/phage synergistic interactions for VRE isolates regardless of the antibiotic class used. Much greater study will be necessary to uncover the basis by which the synergy occurs between these separate antibiotics and phages.

**Figure 6.**
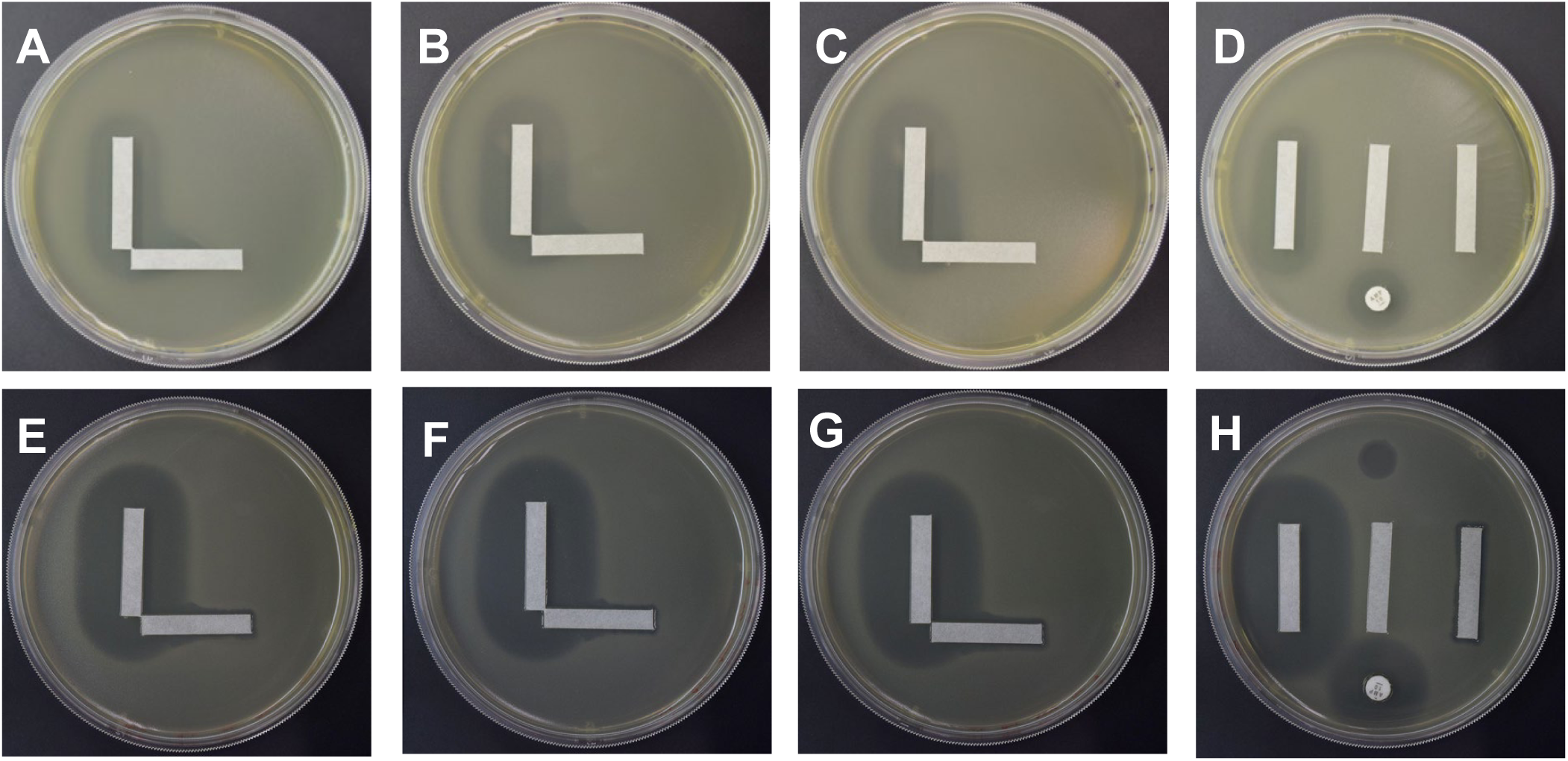
Solid media cooperativity assays for Vancomycin Resistant Enterococcus (VRE). Each specimen was tested with ampicillin (vertical strip) and a bacteriophage (horizontal strip). *E. faecium* NYU with phage Bob is demonstrated in Panels A-D, where A-C represent 3 separate replicates of the cooperativity assay, and panel D represents the control plate with a vertical ampicillin strip (left), blank strip (middle), and phage strip (right), antibiotic disk (bottom), and phage spot (top). *E. faecalis* Yi-6 with phage Bop is demonstrated in Panels E-H, where Panels E-G represent separate replicates and Panel H represents the control plate.

We performed cooperativity assays for a number of different VRE and VSE isolates of *E. faecium* and *E. faecalis*. These assays were performed using antibiotics ampicillin, vancomycin, and linezolid, but also were performed with different myovirus and siphovirus phages infectious for *Enterococcus* spp. We identified a number of isolates in which no evidence of cooperativity could be identified (**Figure S5**). For example, no interactions could be identified for *E. faecium* strain EF208PII nor *E. faecalis* EF140PII. However, there were significant interactions identified for *E. faecium* isolates, including EF98PII, and NYU (**Table S9**), but also for *E. faecalis* strains V587, EF116PII, Yi-6, and B3286. In all our analyses of the patterns of interactions between antibiotics and phages, we did not observe any that matched the models of additivity nor antagonism.

### Evaluation of cooperativity in gram-negative *Stenotrophomonas maltophilia* (STM)

We also analyzed a gram-negative bacterium to identify whether we could observe the same type of synergy that we observed in Enterococcus between antibiotics and phages. We chose the gram-negative bacterium STM because of its profiles of antibiotic resistance, where treatment is often limited to a few antibiotics, including ceftazidime, levofloxacin, and trimethoprim/sulfamethoxazole (**Table S6**) (25). We first tested ceftazidime along with phage KB824 in our cooperativity assay (**Figure 7, Panels A-D**). We identified substantial evidence of synergistic interactions in all replicates tested. We also noted this type of synergistic interaction extended to additional STM strains B28S (**Panels E-H**) and K279a (**Table S9**). We also tested several different phages which were active against our group of STM isolates (**Table S10**). The synergy results were not phage specific, as we identified synergistic interactions for a podovirus (KB824) and a siphovirus (ANB28). However, in experiments using the antibiotic levofloxacin, none of the STM isolates demonstrated evidence of cooperativity with phages (**Figure S6** and **Table S11**). In summary, while we identified some instances of synergistic interactions between ceftazidime and different phages, most of our STM isolates did not show any evidence of cooperativity between antibiotic and phage.

**Figure 7.**
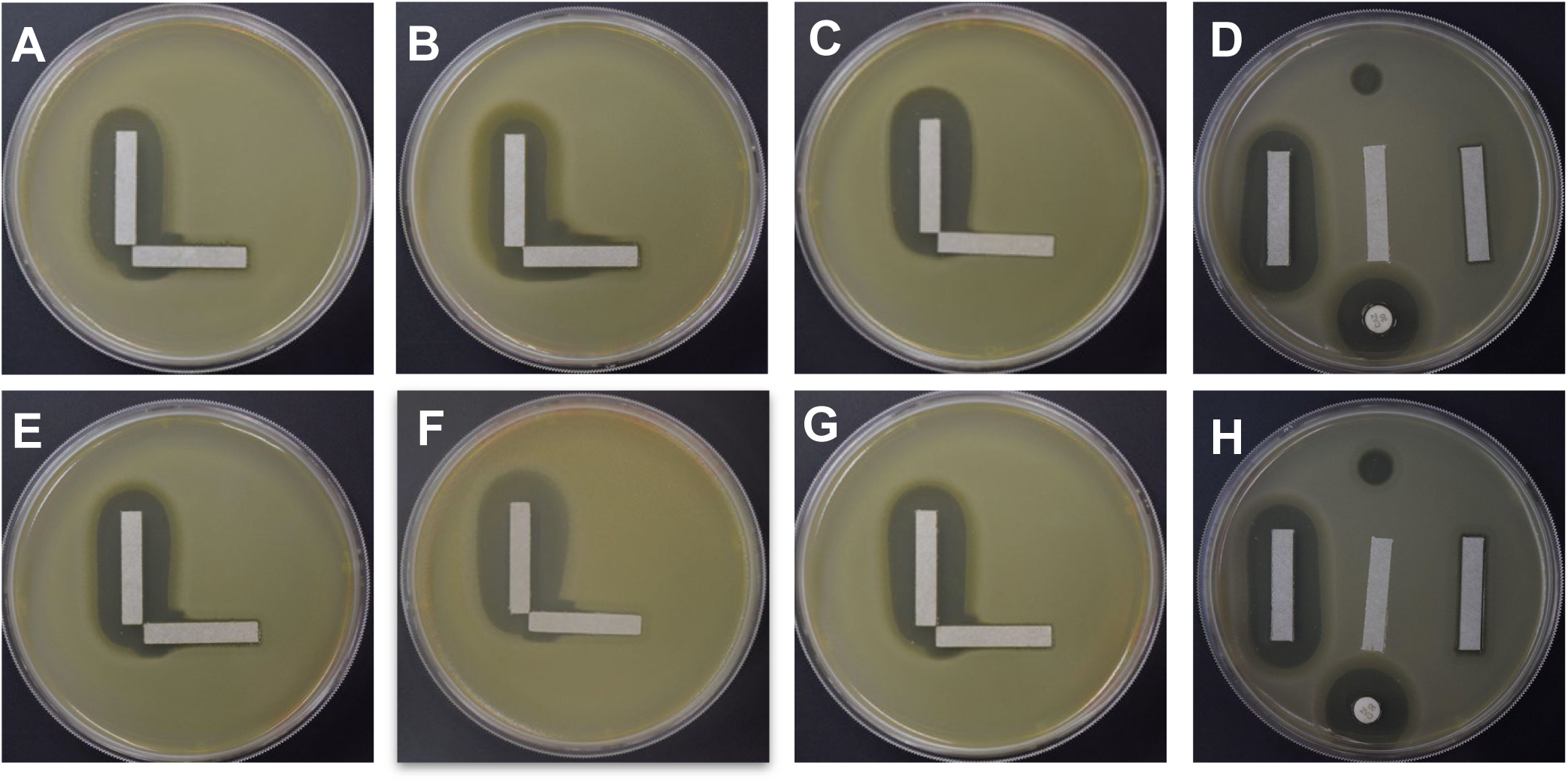
Solid media cooperativity assays for *Stenotrophomonas maltophilia* (STM). Each specimen was tested with ceftazidime (vertical strip) and a bacteriophage (horizontal strip). STM B28B with phage KB824 is demonstrated in Panels A-C, which represents 3 separate replicates of the cooperativity assay. Panel D represents the control plate with a vertical ceftazidime strip (left), blank strip (middle), and phage strip (right), antibiotic disk (bottom), and phage KB824 spot (top). STM B28S with phage KB824 is demonstrated in Panels E-G, which represents separate replicates. Panel H represents the control plate with a vertical ceftazidime strip (left), blank strip (middle), and phage strip (right), antibiotic disk (bottom), and phage KB824 spot (top).

## DISCUSSION

Cooperativity between antibiotics and phages can be difficult to measure and has only recently started to garner greater attention (26-28). In its current state, phages are most often administered concurrently with standard-of-care antibiotics to patients with ARB infections under single patient Investigational New Drug Applications. Because of concurrent antibiotic use, it often is difficult to discern whether the antibiotics, the phage, or the combination of both resulted in improvement. There have been anecdotal cases that demonstrate the potential for cooperative interactions between antibiotics and phages (12, 29), and sophisticated laboratory methods for synergy testing in broth (9), but there are no standardized techniques by which cooperativity is measured. Furthermore, synergy for antimicrobials is generally performed in clinical microbiology facilities (30). Liquid media synergy assays are too complicated to be performed routinely in most clinical laboratories. We developed this solid media cooperativity assay because its simplicity may allow for it to be used broadly across clinical microbiology facilities. While there may be more precise methods, we could develop for characterizing cooperative interactions between antibiotics and phages, the simplicity of the assay we have developed could allow for its adoption across laboratories without the need for expensive equipment.

It was important in the development of this solid media cooperativity assay that we formulate a process that can work for a wide variety of microbes, including gram-positive and gram-negative organisms. There is already a body of literature that suggests such cooperativity, at least in liquid media, may occur (9). In the validation of this assay, we chose to focus on VRE isolates because prior studies have suggested that cooperative interactions can be observed (31). Our data extends those findings to solid medium. The antibiotic resistant nature of VRE makes it an ideal candidate for our analysis because it can cause deep and long-lasting infections that require alternative therapies such as phages (32). We also evaluated STM as an example of a gram-negative organism, as its antibiotic resistant nature significantly limits antibiotic treatment options (33). STM also is capable of causing long-lasting infections due to its ability to infect those in the Cystic Fibrosis population, where the organism can be incredibly difficult to eradicate (34). Our finding of synergistic interactions between phages and the antibiotic ceftazidime may restore the ability to use this antibiotic for these STM infections, where we observed synergy largely in STM isolates that showed intermediate MICs to ceftazidime alone. Future work will be necessary to determine how the phage may restore the susceptibility to ceftazidime, and it may be through reduced expression or efficacy of the L1 and L2 beta-lactamases, or via changes in cell wall composition in response to the phage. We hypothesize that synergistic interactions between antibiotics and phages are not limited to the Enterococcus and STM isolates used in this study but can likely be extended to further ARB such as the ESKAPE pathogens that are often the target of phage therapies.

Identifying synergistic interactions in an *in vitro* study such as this does not necessarily predict what may occur when such treatments are utilized *in vivo*. However, prior studies have indicated that *in vitro* responses may predict the utility of such treatments in humans (35). Even though antibiotics and phages are used together in the majority of phage therapy clinical cases, the combination has been understudied to date (36). We hope to alter this standard approach by implementing an easy to perform assay for identifying phage-antibiotic synergy. Thus, as an increasing number of phage therapy cases take place, physicians can be provided with information to better inform their decisions on whether antibiotics and phages may have cooperative effects.

Anecdotal studies indicate that the administration of both vancomycin and phages may have synergistic activity against VRE (31). While the mechanisms behind such interactions have not been well studied, our data help to confirm those findings and extend them to an easy to perform solid media assay. The currently used broth-based assays are cumbersome and require specific equipment which makes widespread adoption in clinical laboratories difficult.

We show that there are synergistic interactions between vancomycin and phages with myovirus and siphovirus morphologies (37) from for both *E. faecalis* and *E. faecium*. While we are not aware of specific instances where clinical treatments have taken place for VRE isolates using vancomycin and phages, the *in vitro* data shown here suggests that there is the potential for clinical efficacy. One of the simplest clinical rules available for the treatment of VRE has been to avoid the use of vancomycin (38). Our confirmation of the finding that vancomycin in combination with phages may restore the utility of vancomycin in the treatment of VRE could be of significant benefit in the treatment of this life-threatening pathogen. We identified synergistic interactions for other antibiotics, including ampicillin and linezolid (**Figures 5 and 6**), which suggests that a broad array of antibiotics may be available for treatment of VRE when phages are involved, even in cases where the VRE isolates are initially resistant to the antibiotics.

There is a lack of standardization of techniques by which to deliver phage therapy and to choose which antibiotic/phage combinations may be the most efficacious (39). We developed the solid media cooperativity assay presented here with the goal to help standardize techniques for decision-making in phage therapy cases and to allow for a much wider adoption of techniques for identifying cooperativity between antibiotics and phages. Our results indicate that this assay is robust and reproducible, can be extended to both gram-positive and gram-negative bacteria, can be applicable across different phage morphologies, applies to multiple different antibiotics, and does not necessarily require pre-existing antibiotic nor phage susceptibility in the target bacteria for cooperativity to be observed. We believe solid media assays for the detection of phage/antibiotic cooperativity should serve as standard adjunctive testing to help guide the use of antibiotics and phages in phage therapy cases.

## Abbreviations

STM: *Stenotrophomonas maltophilia*
VRE: Vancomycin Resistant Enterococcus
VSE: Vancomycin Susceptible Enterococcus
ARB: Antibiotic Resistant Bacteria
ESKAPE: *Enterococcus faecium*, *Staphylococcus aureus*, *Klebsiella pneumoniae*, *Acinetobacter baumannii*, *Pseudomonas aeruginosa*, and *Enterobacter sp*.
MBC: Minimum Bactericidal Concentration

## Author Contributions

Conceived and designed project: DTP, KW, RL, SIF, AM, SD, and AR.

Performed experiments: EK, JO, JMJ, ANB, and AR.

Analyzed the data: EK, JO, JMJ, AG, SD, AM, PG, KW, DTP, and SIF.

Wrote and/or edited the manuscript: DTP, KW, AGCG, EK, JO, AK, RS, SA and SIF.

Provided materials for the study: MC, PK, and MP.

All authors have reviewed the manuscript.

## Acknowledgements

We thank the UCSD Clinical Microbiology Laboratory for their participation in this work.

## Figure Legends

**Figure S1:** Plate configurations for screening of antibiotic-phage cooperativity. The test plate (Panel A) and control plate (Panel B) configurations are shown.

**Figure S2.** Possible patterns to be observed for phage-antibiotic cooperativity assays. Cooperativity (Panel A), and no cooperativity (Panel B) are shown.

**Figure S3:** Whatman filter strips loaded with antibiotic or bacteriophage solutions are placed perpendicularly on an agar plate that has an *E. faecium* bacterial lawn. As the antibiotic and phage solutions diffuse through the agar, they interact with bacteria, killing the bacteria in regions where an effective minimum bactericidal concentration (MBC) is reached. The interface between live and dead bacteria creates a profile that aligns with the MBC for the antibiotic, phage, and bacteria combination. A computational model can be used to predict the concentrations of the antibiotic and bacteriophage solutions as they diffuse through the agar. The effective combinatory concentration can be calculated by making assumptions about the antibiotic and phage interactions (*i.e.* no interaction, additive, synergistic, antagonistic).

**Figure S4:** Stamping Procedure. 1. Dried bacteriophage and antibiotic strips are aligned at 90° using the right-angle edge of the stamp column. The square region marked by arrow indicates that strips are not overlapping and aligned at 90°. 2. The solidified plate is inversed and gently stamped onto the aligned strips. 3. Top view of the stamping process. L-shape should be stamped so that there is plenty of room for bacteriophage and antibiotic strips to demonstrate proper clearing.

**Figure S5.** Solid media cooperativity assays for Vancomycin Resistant Enterococcus (VRE). *E. faecium* EF208PII with antibiotic vancomycin and phage PL is represented in Panels A-D. *E. faecium* EF98PII with phage Ben and antibiotic ampicillin is represented in Panels E-H. *E. faecium* EF208PII with phage PL and antibiotic linezolid is represented in Panels I-L. Each specimen was tested with an antibiotic (vertical strip) and a bacteriophage (horizontal strip). Panels D, H, and L represent the control plate with a vertical antibiotic strip (left), blank strip (middle), phage strip (right), antibiotic disk (bottom), and phage spot (top).

**Figure S6:** Solid media cooperativity assays for *Stenotrophomonas maltophilia* (STM). Each specimen was tested with levofloxacin (vertical strip) and a bacteriophage (horizontal strip). STM SM17 with phage 2φ2 is demonstrated in Panels A-D, where A-C represent 3 separate replicates of the cooperativity assay, and panel D represents the control plate with a vertical levofloxacin strip (left), blank strip (middle), phage strip (right), antibiotic disk (bottom), and phage spot (top). STM SM26 with phage KB824 is demonstrated in Panels E-H, where Panels E-G represent separate replicates and Panel H represents the control plate. STM SM27 with phage ANB28 is represented in Panels I-L.

